# *‘Candidatus* Megaira’ are diverse symbionts of algae and ciliates with the potential for defensive symbiosis

**DOI:** 10.1101/2022.09.23.509160

**Authors:** Helen R. Davison, Gregory D. D. Hurst, Stefanos Siozios

## Abstract

Symbiotic microbes from the genus ‘*Candidatus* Megaira’ (Rickettsiales) are known to be common associates of algae and ciliates. However genomic resources for these bacteria are scarce, limiting our understanding of their diversity and biology. We therefore utilized SRA and metagenomic assemblies to explore the diversity of this genus. We successfully extracted four draft ‘*Ca*. Megaira’ genomes including one complete scaffold for a ‘*Ca*. Megaira’ and identified an additional 14 draft genomes from uncategorised environmental Metagenome-Assembled Genomes. We use this information to resolve the phylogeny for the hyper-diverse ‘*Ca*. Megaira’, with hosts broadly spanning ciliates, micro- and macro-algae, and find that the current single genus designation ‘*Ca*. Megaira’ significantly underestimates their diversity. We also evaluate the metabolic potential and diversity of ‘*Ca*. Megaira’ from this new genomic data and find no clear evidence of nutritional symbiosis. In contrast, we hypothesize a potential for defensive symbiosis in ‘*Ca*. Megaira’. Intriguingly, one symbiont genome revealed a proliferation of ORFs with ankyrin, tetratricopeptide and Leucine rich repeats like those observed in the genus *Wolbachia* where they are considered important for host-symbiont protein-protein interactions. Onward research should investigate the phenotypic interactions between ‘*Ca*. Megaira’ and their various potential hosts, including the economically important *Nemacystus decipiens*, and target acquisition of genomic information to reflect the diversity of this massively variable group.

**Data Summary:** Genomes assembled in this project have been deposited in bioproject PRJNA867165

**Impact statement:** Bacteria that live inside larger organisms commonly form symbiotic relationships that impact the host’s biology in fundamental ways, such as improving defences against natural enemies or altering host reproduction. Certain groups like ciliates and algae are known to host symbiotic bacteria commonly, but our knowledge of their symbiont’s evolution and function is limited. One such bacteria is ‘*Candidatus* Megaira’, a Rickettsiales that was first identified in ciliates, then later in algae. To improve the available data for this common but understudied group, we searched the genomes of potential hosts on online databases for Rickettsiales and assembled their genomes. We found 4 ‘*Ca*. Megaira’ this way and then used these to find a further 14 genomes in environmental metagenomic data. Overall, we increased the number of known ‘*Ca*. Megaira’ draft genomes from 2 to 20. These new genomes show us that ‘*Ca*. Megaira’ is far more diverse than previously thought and that it is potentially involved in defensive symbioses. In addition, one genome shows striking resemblance to well characterized symbiont, *Wolbachia*, in encoding many proteins predicted to interact directly with host proteins. The genomes we have identified and examined here provide baseline resources for future work investigating the real-world interactions between the hyper diverse ‘*Ca*. Megaira’ and its various potential hosts, like the economically important *Nemacystus decipiens*.

## Introduction

A wide range of bacteria species reside as endosymbionts in both microeukaryotes, and algae [1–8]. Symbiont presence can affect the biology of their host in significant ways, from reproductive manipulation [9, 10] to stress tolerance [11], nutrient production [12, 13] and methanogenesis [14]. Symbionts in microeukaryotes were recognised as early as 1902 in the amoeba *Pelomyxa* [15]. Whilst some systems are well understood, such as *Caedibacter* and *Paracaedibacter* in *Paramecium* [10], our knowledge of symbiont evolution and function in microeukaryotes is fragmented in comparison to symbioses in animals and terrestrial plants. For instance, the effects of endosymbiotic bacteria in algae are currently unknown, with studies rarely extending beyond the presence of the symbioses and the phylogenetic affiliation of the symbiont [7, 8, 16].

In the last decade Rickettsiales have been identified as a group that commonly form symbioses with microeukaryotes as well as invertebrates and algae [4, 17–23]. The origins of some families within the Rickettsiales, like the Rickettsiaceae, may derive from symbioses with microeukaryotes [20]. ‘*Ca*. Megaira’ is a member of Rickettsiales and a relative of *Rickettsia* and *Wolbachia* which are prolific endosymbionts with wide ranging effects on their hosts REF. As such, ‘*Ca*. Megaira’ has the potential to impact its hosts in many ways. However, few functional studies have been completed, which found that ‘*Ca*. Megaira’ presence improved growth in some *Paramecium* [19, 24]. However, in contrast to *Rickettsia* and ‘*Ca*. Tisiphia’, there is currently very limited genomic data for ‘*Ca*. Megaira’, with a single closed and a single draft genome, both from algae [25].

The increasing power and reliability of bioinformatic tools now enable us to extract high quality microbial symbiont genomes from the Sequence Read Archive (SRA) deposits [25, 26]. We can search for symbiotic bacteria in hosts without *a priori* hypothesis to establish novel symbiotic interactions with target microbes, and then assemble draft genome sequences for the symbionts. Declining costs have driven a surge in sequencing non-model taxa like microeukaryotes, providing ample data from which to extract symbiont genomic data. For taxa like ‘*Ca*. Megaira’ where there is little genomic information available, this data then provides us with the opportunity to explore their evolution and diversity in more detail and generate hypotheses as to the function of the symbioses found.

In this study, we search and extract potential ‘*Ca*. Megaira’ symbionts in GenBank SRA data for ciliates and all current classifications of micro- and macro-algae. In addition, we identified ‘*Ca*. Megaira’ genomes amongst publicly available Metagenome-Assembled Genomes (MAGs) in GenBank. These data collectively expand the known whole genomes of ‘*Ca*. Megaira’ from two to twenty genomes and enable phylogenomic and metabolic analyses.

## Methods

### Collection of external genomes for metagenomics and phylogenomics

Illumina SRA data for all ciliates and current classifications of Algae as of 05 May 2021 were downloaded from NCBI to screen for symbiont genomes. These were: Bacillariophyceae, Charophyceae, Chlorarachniophytes, Chlorophyceae, Chlorophyta, Chrysophyceae, Cillophora, Cryptophyceae, Dictyochophyceae, Dinophyceae, Euglenophyceae, Eustigmatophyceae, Haptophyta, Mesostigmatophyceae, Phaeophyceae, Rhodophyta, Synurophyceae, Ulvophyceae, Ulvophyceae. Libraries were excluded if they were: Extremely shallow sequencing efforts below 500 megabases, macronucleus-only sequencing, mutant resequencing, listed as antibiotic treated, or dd-RAD sequence. In total 1113 of 3445 algae and 464 of 547 ciliate libraries were identified for onward analysis.

### Metagenomic identification, assembly of genomes and phylogenomic analysis

SRA deposits were screened for the presence of Rickettsiales using Phyloflash [27]. Rickettsiales-positive libraries were taken forward for metagenomic assembly and binning to extract full genome sequences as described in Davison *et al*. (2022) [25]. Briefly, metagenomic assembly, binning and quality check was performed with Megahit, Metabat2 and CheckM [25, 28–31]. Samples that contained >50% complete symbiont genomes with <5% contamination were taken forward for further examination and manual refinement. GTDBtk [32] was used for taxonomic classification of each extracted genome and identify their nearest relatives. Genome bins identified as Rickettsiales were named as follows: first three letters of their closest relative + first letter of host genus + first four letters of host species + bin number. For example, a ‘*Ca*. Megaira’ from a *Mesostigma viride* SRA in bin 4 would be labelled MegMviri4. *Nemacystus decipiens* (bioproject PRJDB7493) had multiple SRA libraries from the same biosample which we co-assembled with Megahit. Then, each library was individually mapped back to the assembly with bowtie2 [33] and symbiont bins were identified with Metabat2. Nine of the libraries were mate-pair reads with insert sizes ranges from 2-13 kbp and were used to scaffold the draft assembly and close the genome using BESST algorithm [34].

Additional ‘*Ca*. Megaira’ genomes were identified on GenBank by cross examining core protein sequences of our new draft genomes with the non-redundant protein sequence database through Blastp [35]. The method identified 14 existing, unclassified environmental MAGs related to known ‘*Ca*. Megaira’.

In order to anchor our genomes against previous knowledge of ‘*Ca*. Megaira’ diversity, 16S rRNA sequences were assembled for ‘*Ca*. Megaira’ symbionts where possible, though due to the limitations of metagenomic binning and assembly this was not possible for several environmental metagenomes. MegHsini1 is a partial genome and two 16S rRNA sequences can be extracted with Anvi’o 7 [36]. The most complete of these was used for 16S rRNA sequence placement. The less complete one seems to be related to *Deineraceae* and was deemed a likely contaminant. Additional sequences can be found in Supplementary table S7.

The draft genome data were used to enable a phylogenomic approach to ‘*Ca*. Megaira’ diversity alongside existing known ‘*Ca*.Megaira’ genomes (Supplementary Tables S1 and S2). Orthologous genes across the 20 Megaira genomes were identified using Anvi’o 7 [36] for the purpose of extracting the core gene clusters (50 gene clusters). Average Nucleotide Identity (ANI) was calculated through pyANI within Anvi’o 7 (Supplementary Table S4). Average Amino-acid Identity (AAI) was calculated pairwise for each genome pair through the AAI-Matrix calculator from the enveomics toolbox (Supplementary Table S3) [37]. Synteny between JAFLDA01 and MegNEIS296 was established with PROmer in MUMmer3 package [38]. Maximum likelihood trees were produced with IQ-Tree and automatic best model selection using ModelFinder [39, 40] with 1000 replicates of UltraFast Boostrap [41] and SH-like Approximate Likelihood Ratio Test [42]. Models selected for each tree are as follows: Megaira core amino acids = LG+F+I+G4, Megaira 16S rRNA = GTR+F+R3.

### Examining metabolic potential, annotation and identifying NRPS systems

High quality genomes from the above were defined as >90% complete and contamination <10%. This process defined 2 existing ‘*Ca*. Megaira’ genomes (MegCarteria, and MegNEIS296), 3 novel genomes derived from the SRA (MegSroe9, MegMviri4, and MegNdeciBESST), and 5 novel genomes derived from MAGs (JAFLDA01, VGEX01, JAJTEJ01, NVVL01, JAFLCZ01) as high quality, and these were analysed alongside a ‘*Ca*. Tisiphia’ genome and *Orientia*. Metabolic potential was predicted based on KEGG annotations by Anvi’o 7 [36, 43]. Heatmaps of pathway completeness were sorted by phylogeny and plotted in Python with Seaborn [44, 45]. An upsetplot of shared gene clusters between genomes was constructed with ComplexUpset [46] in R 4.1.0 [47].

AntiSMASH [48] was then used on the eight high quality genomes to predict secondary metabolites such as those produced by the non-ribosomal peptide synthetase (NRPS) systems. These have been identified previously in the existing Megaira genome, MegNEIS296 (ASM2041082v1). Clinker was used to visualise the similarity between the resulting systems found [49]. Further annotations were made with InterProScan 5 [50]using Pfam, TIGRFAM, PANTHER and GOterms.

## Results

### Assembly of genomes

After metagenomic binning, 4 SRA deposits were identified as harbouring ‘*Ca*. Megaira’ and taken forward for further analysis. All but one genome is >90% complete (Table x). MegHsini1 is derived from a single cell genomics approach and was just 62.84% complete and thus not included in onward metabolic analyses. However, MegHsini1 core genes clusters and marker genes were retained for phylogenetic placement. No Rickettsiales other than ‘*Ca*. Megaira’ were recovered. The ‘*Ca*. Megaira’ from *Nemacystus decipiens* (PRJDB7493) was the only genome that could be assembled into one scaffold, albeit not closed, using the available matepair data. This genome, named here as MegNdeciBESST, MegNdeciBESST, has a total size of about 1.3Mb and contains 20 gaps, ranging from 346 to 5679 bp. 14 additional environmental MAGs, previously characterized as unclassified Rickettsiales, were identified in GenBank. These environmental MAGs are of similar quality as the MAGs constructed from SRA databases here (Table 1).

**Table 1.**
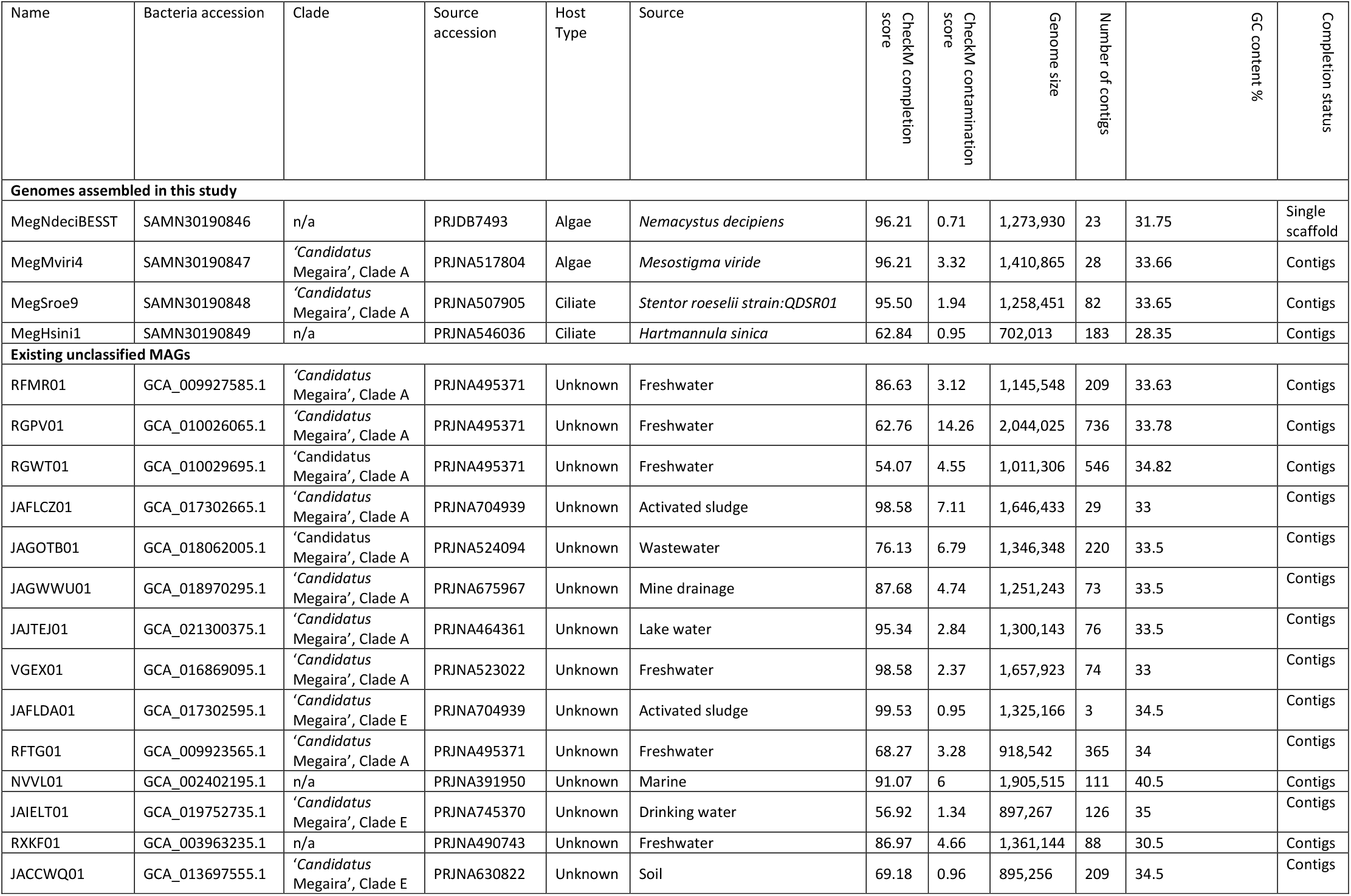
Genome statistics and sources, in depth metadata including SRA sample accessions can be found in the supplementary.

### Phylogeny and evolution

ANI and AAI scores alongside phylogenetic analysis suggest that the whole of ‘*Ca*. Megaira’ clade is deeply divergent (Figures 1–4, and Supplementary figure 1). For instance, AAI scores between Clade A and Clade E are <65% (Figure 3), and there is no synteny between MegNEIS296 and JAFLDA01, representatives of each group (supplementary figure 2). The existing ‘*Ca*. Megaira’ clades do not sufficiently describe the diversity seen within the group and our genomic data suggest that the ‘*Ca*. Megaira’ clade groups may represent different genera.

**Figure 1.**
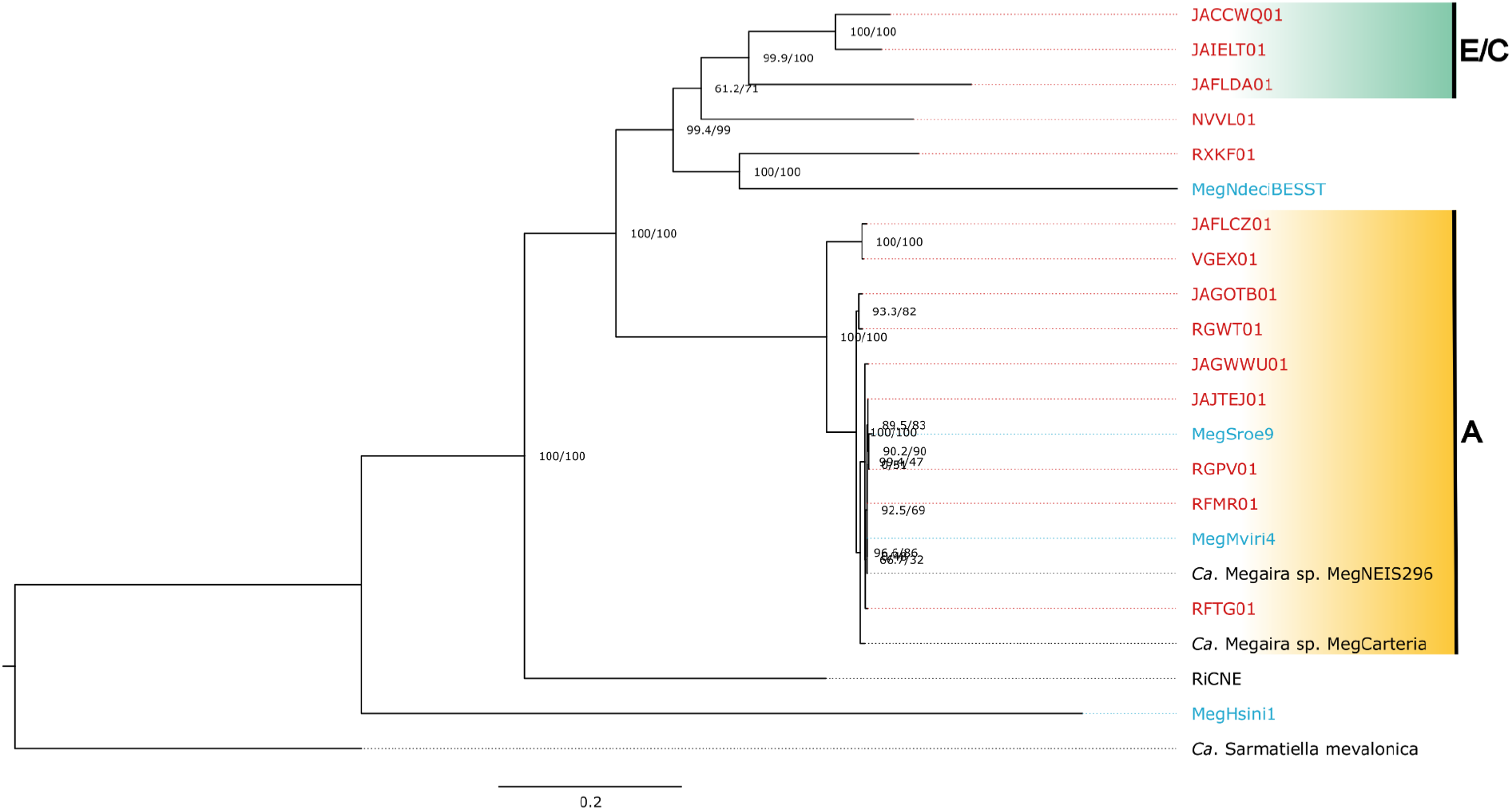
Ca. Megaira core genome maximum likelihood tree with 1000 SH-aLRT and ultrafast bootstrap (UFB) support. Support for each split is shown as (SH-aLRT/UFB), with strong support being >=80/>=95 Samples from this study are blue and existing environmental metagenomes are red.

Four of the ‘*Ca*. Megaira’ draft genomes (MegHsini1, MegNdeciBESST, NVVL01, and RXKF01) represent new ‘*Ca*. Megaira’ clades (figure 1 and 2). AAI scores of <65% suggest that these four are sufficiently derived to be considered new genera (Figure 3). However, the placement of MegHsini1 within the Rickettsiales is currently uncertain (Figure 1, 2 and Supplementary figure 1). For instance, GTDBtk classification does not assign MegHsini a genus or species (Supplementary Table S9). Based on available 16S rRNA and supporting AAI scores, most of the MAGs clustered within Clade A; three MAGs fall into Clade E (and possibly clade C); and two form a new group within Clade A which share an ANI similarity score of <95% (Figures 2 and 3). Two MAGs lack 16S rRNA sequence and cannot currently be associated with any group as 16S rRNA is the only marker used to date to classify ‘*Ca*. Megaira’.

**Figure 2.**
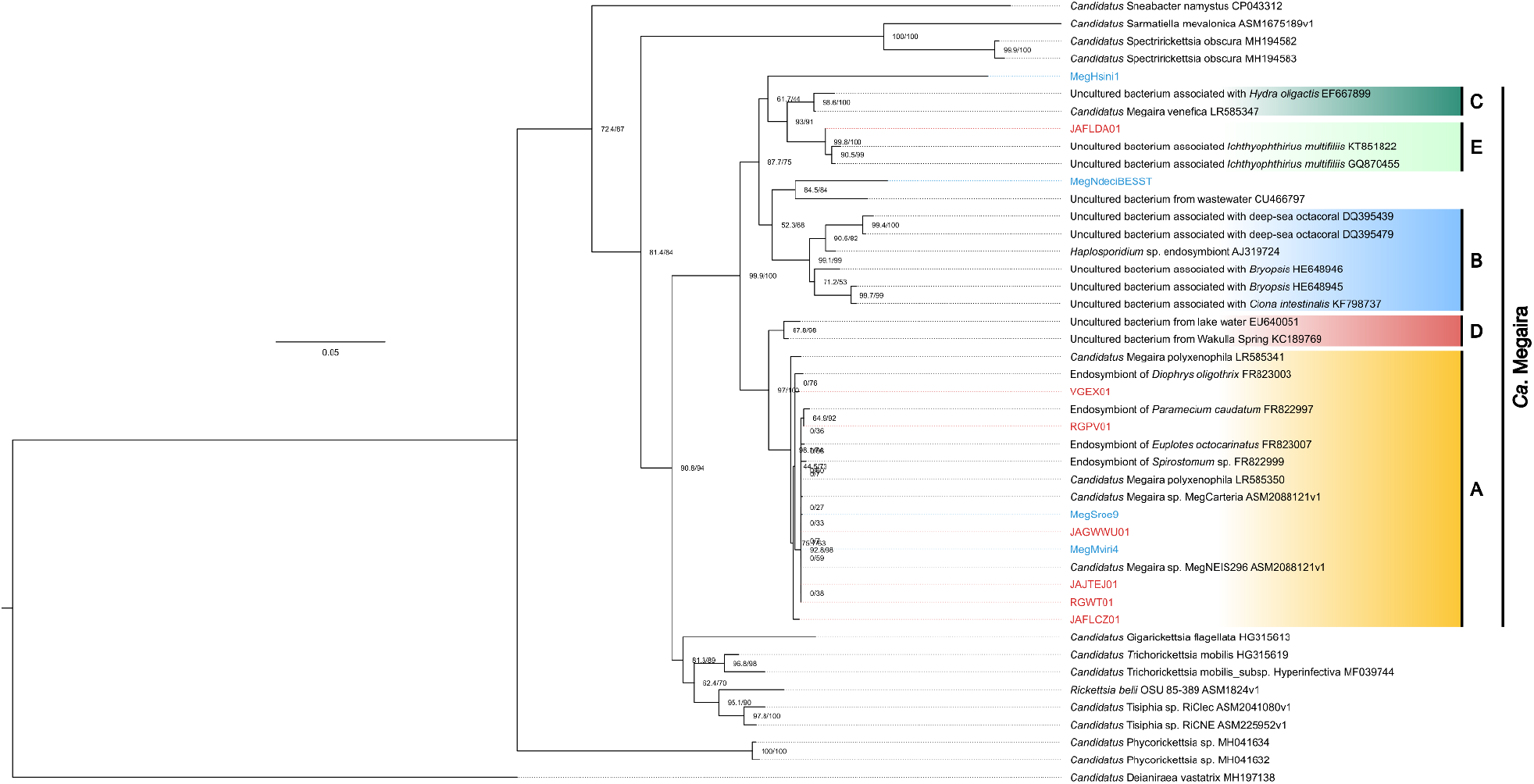
Megaira 16S rRNA maximum likelihood tree with 1000 SH-aLRT and ultrafast bootstrap (UFB) support. Support for each split is shown as (SH-aLRT/UFB), with strong support being >=80/>=95. Samples from this study are blue and existing environmental metagenomes are red.

In several instances the genomes used in this study are the only ones available for their lineage (Figure 1). In addition, MegHsini1 is very incomplete (62%) in comparison to the majority others (11 of 18 are >85% complete, Table 1), despite having high depth of coverage (~245X, Supplementary table S1). Although a 16S rRNA sequence was also recovered, MegHsini1 also is weakly placed in its phylogenies (Figure 1 and 2, and Supplementary figure 1) and potentially suffers from long branch attraction. At this stage we do not know if MegHsini1’s uniqueness is a genuine feature, or a symptom of fragmentation caused by amplification bias during the enrichment steps of single cell genomics. Further expansion of genomic data for ‘*Ca*. Megaira’ is required to refine the phylogeny of the bacteria in this species, and we would recommend any future screening efforts use other indicator genes alongside 16S rRNA. The genomic information obtained here will enable development of these markers and PCR protocols.

**Figure 3.**
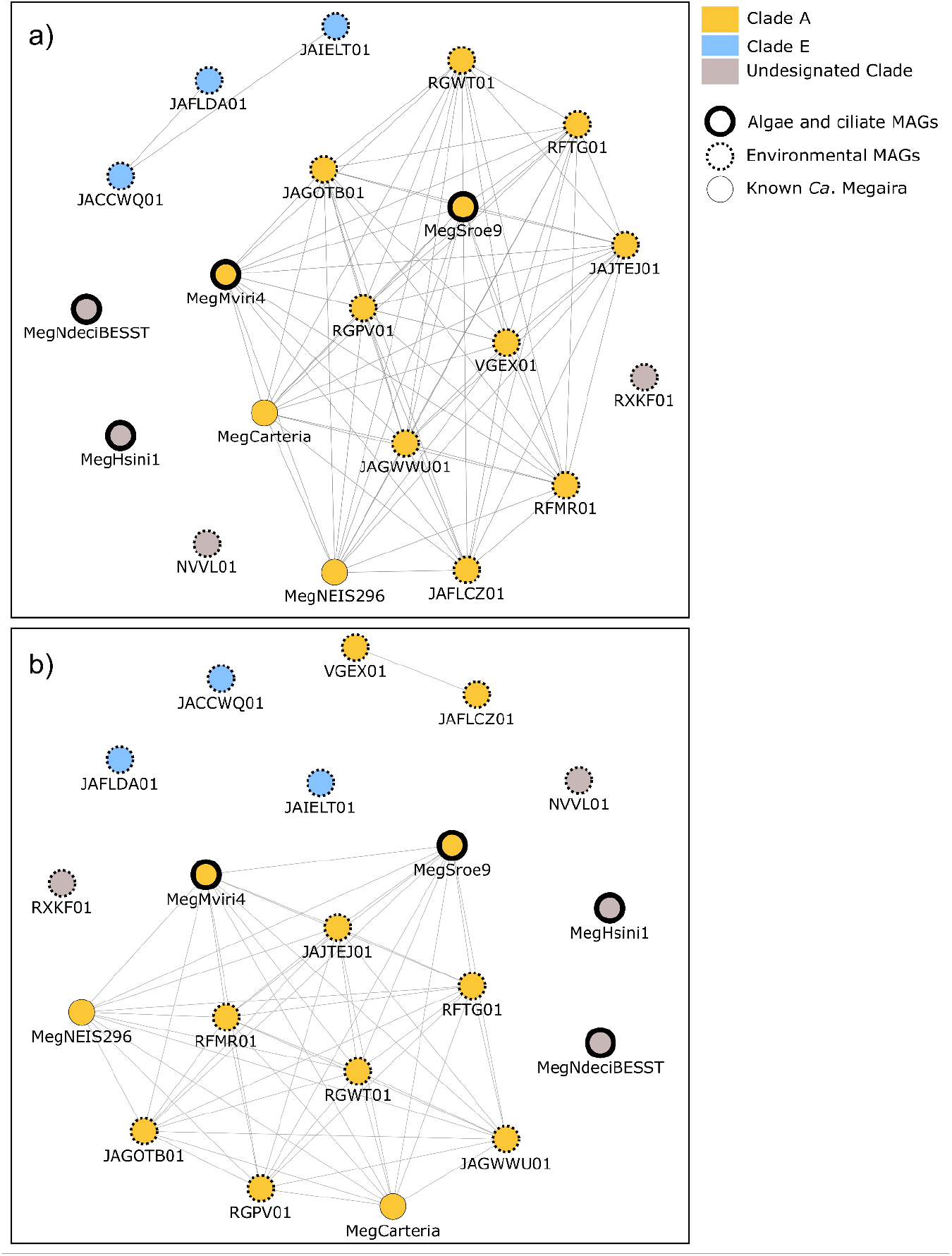
*AAI and ANI map for ‘Ca*. Megaira’ *showing a) genomes sharing >65% AAI similarity and b) genomes with >95% ANI similarity. Raw data can be found in Supplementary Table S3 and S4*.

Gene content analysis across the ‘*Ca*. Megaira’ clades mirror these findings. The A group ‘*Ca*. Megaira’ have a common shared unique gene set and have similar patterns of gene presence absence (Supplementary Figure 1, Figure 4). Outside of clade A stains, NVVL01 is highly distinct, having over double the number of unique gene clusters compared to all other taxa; a large number of unique gene clusters were additionally observed in the other two non-A group strains, MegNdeciBESST, and JAFLDA01 (Figure 4).

**Figure 4.**
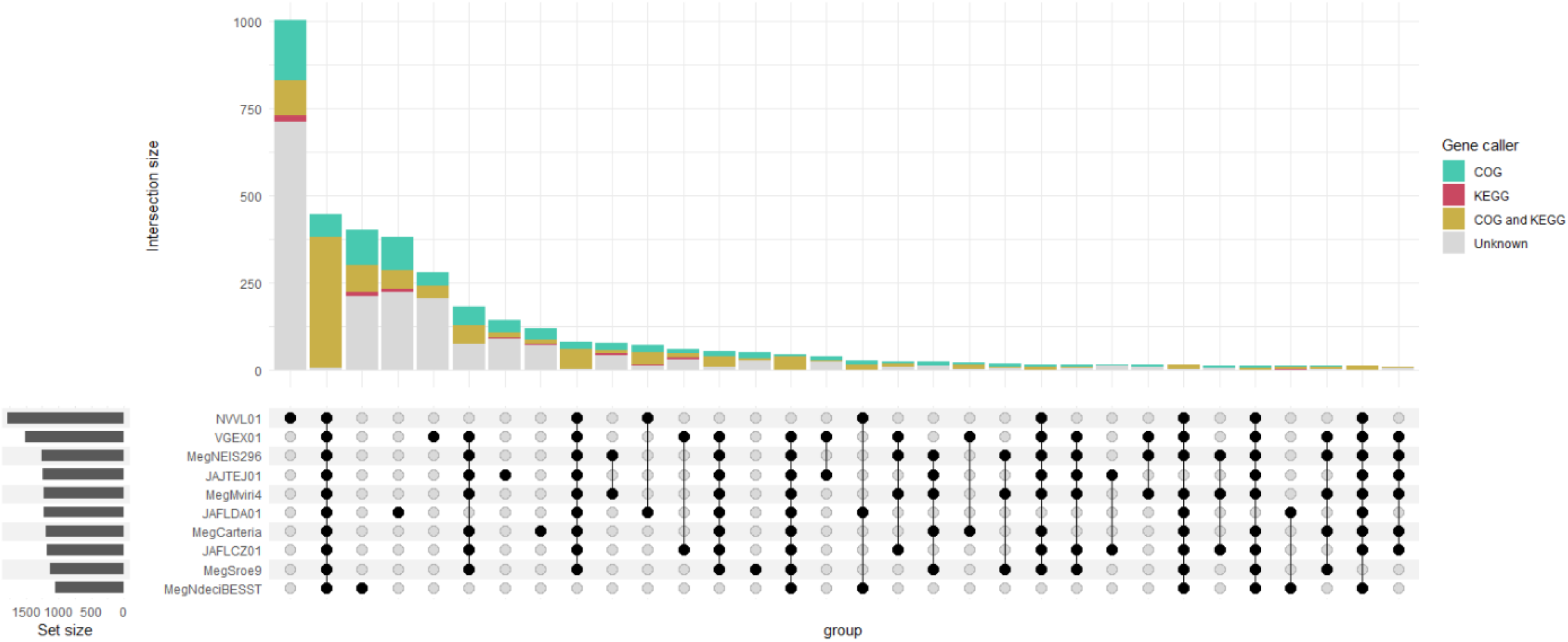
An upset plot showing the number of gene clusters (bars) shared between genomes. Genomes being compared are indicated with black circles and lines. The number of known genes and the caller that identified them are indicated by bar colour. Presence-absence data can be found in Supplementary table S8.

**Supplementary figure 1.**
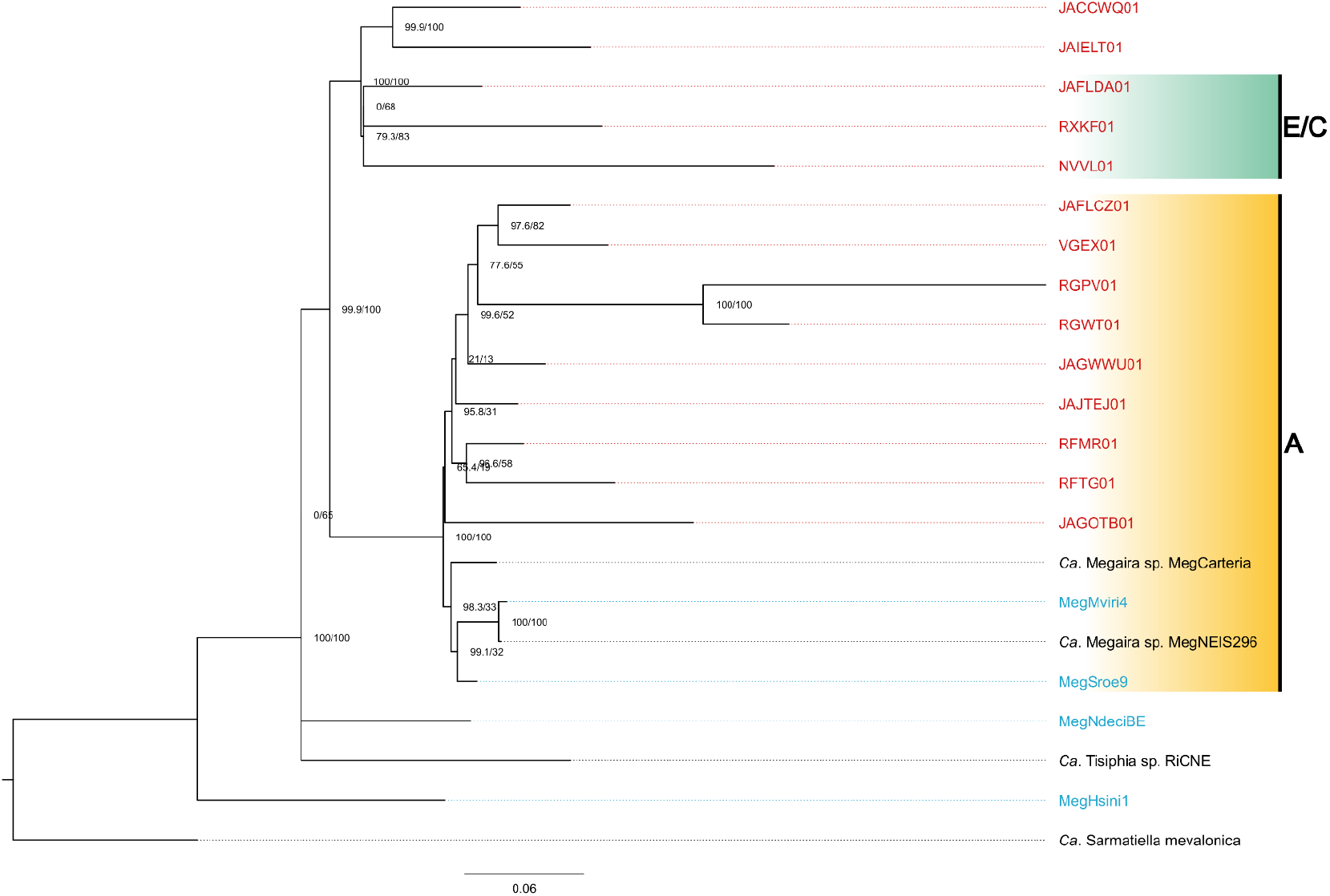
*Maximum likelihood phylogeny of gene cluster presence absence for ‘Ca*. Megaira’ *with 1000 SH-aLRT and ultrafast bootstrap (UFB) support. Support for each split is shown as (SH-aLRT/UFB), with strong support being >=80/>=95 Samples from this study are blue and existing environmental metagenomes are red*.

**Supplementary figure 2.**
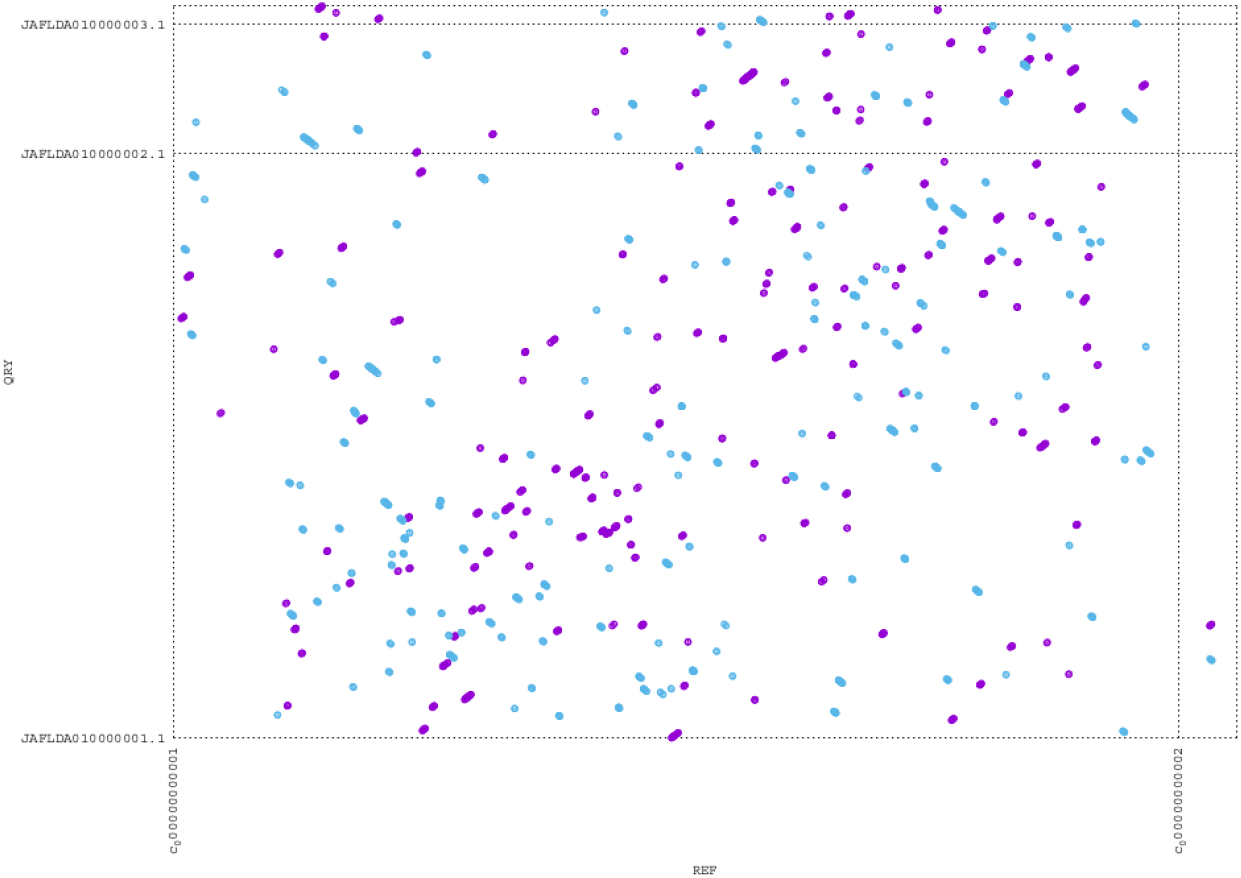
PROmer synteny map constructed by mummer, comparing the similarities in patterns of protein loci across JAFLDA01 and MegNEIS296. Purple are forward matches; blue are reverse matches.

### Metabolism, secondary compound synthesis, secretion systems and potential symbiosis factors

‘*Ca*. Megaira’ are not predicted to encode complete cofactor or vitamins pathways as would be typically observed in nutritional symbioses (Figure 5, Supplementary table S5 and S6). The genome JAFLDA01 is predicted to encode partial thiamine and biotin biosynthesis pathways, which are unlikely to be functional. All ‘*Ca*. Megaira’ are predicted to have complete Nonoxidative Pentose Phosphate Pathways like their relatives, ‘*Ca*. Tisiphia’ (== Torix Group *Rickettsia*) [25, 51]. MegNdeciBESST and JAFLDA01 have complete dTDP-L-Rhamnose pathways (Figure 5). Clade A Megaira, excluding MegCarteria, and clade E Megaira appear to be enriched for terpenoid and polyketide biosynthesis pathways compared to other taxa (Figure 5 and 6).

**Figure 5.**
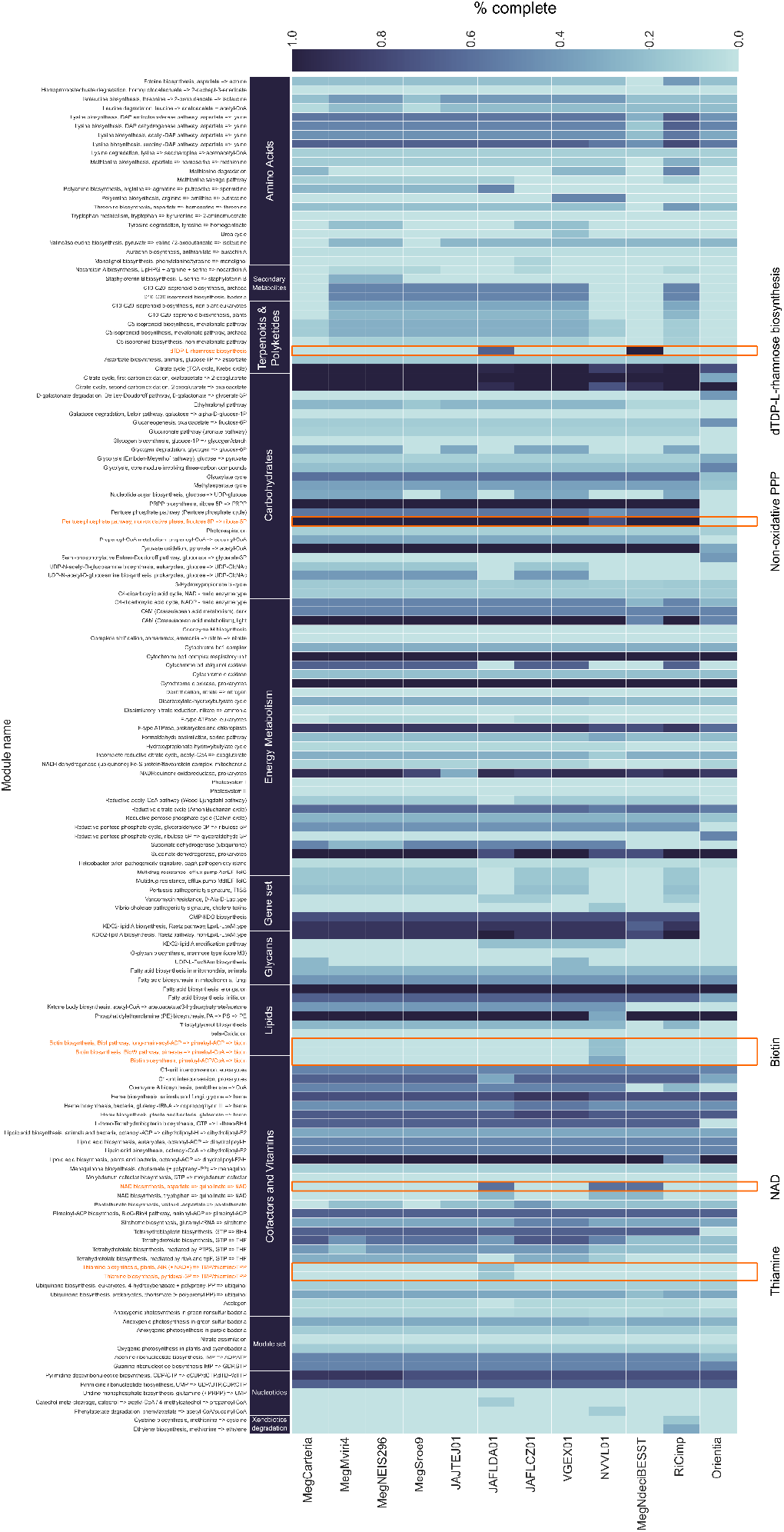
Metabolic heatmap of ‘Ca. Megaira’, with ‘Ca. Tisiphia’ RiCimp and Orientia tsutsugamushi as outgroups. Kofam module completeness from highest to lowest is shown with dark to light blue shading and pathways of interest are highlighted and circled with orange. Full metadata can be found in supplementary table S6.

**Figure 6.**
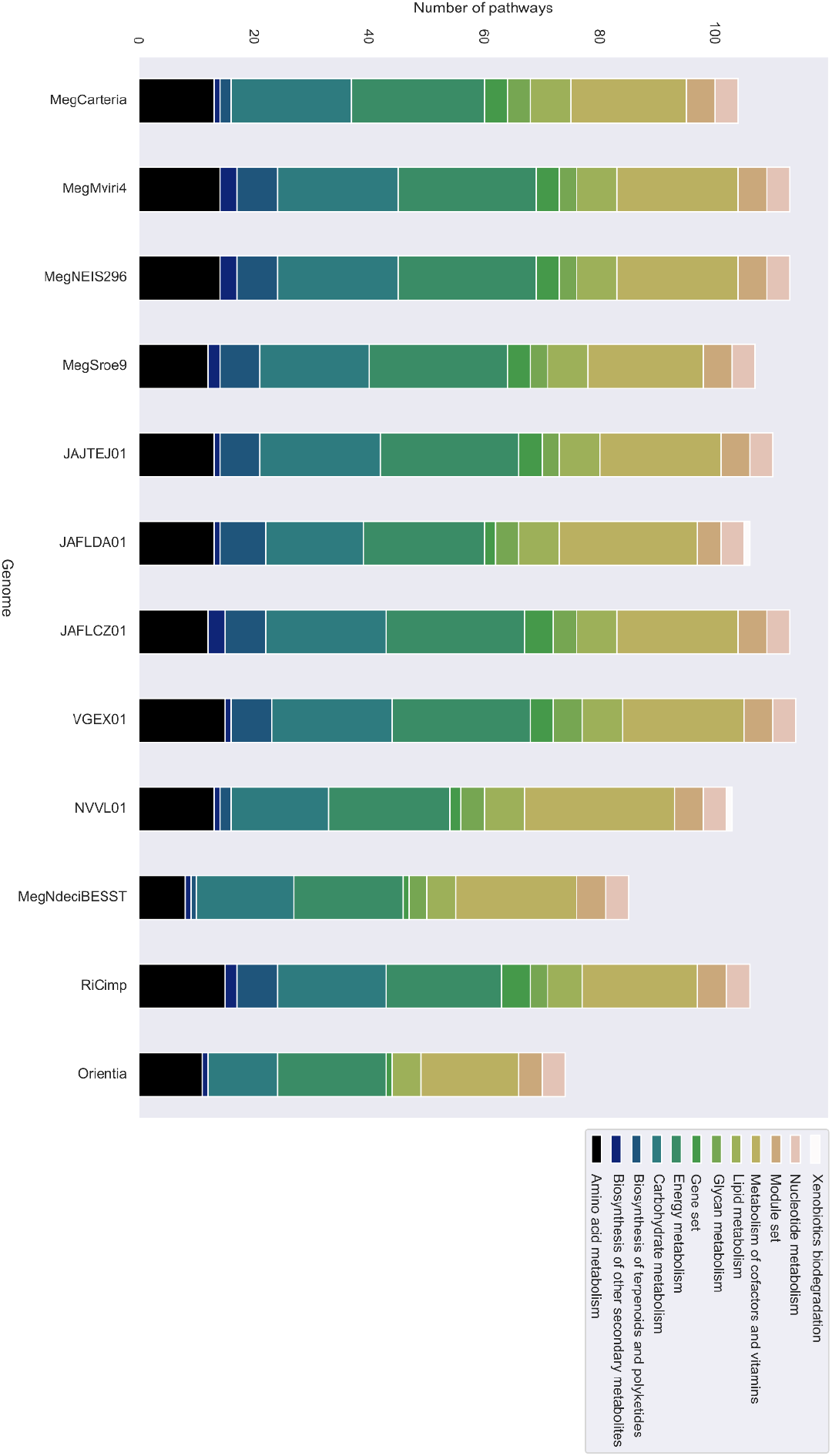
Number of pathways found per genome annotated by KEGG module category for ‘Ca. Megaira’, with ‘Ca. Tisiphia’ RiCimp and Orientia tsutsugamushi as outgroups. Full metadata can be found in supplementary table S5.

AntiSMASH identified five putative Non-Ribosomal Peptide Synthetase (NRPS/PKS) systems in four of eight genomes examined (Figure 7). It also predicted three predicted cyclodipeptide synthases (CDPS), and two ribosomally synthesized and post-translationally modified peptides systems (RiPPs), including one synthesizing a lasso peptide (figure 7, supplementary data). Blastp found that the MegMviri4 contig containing the putative NRPS has 100% similarity with the NRPS found previously in MegNEIS296, albeit it is only a partial fragment. Considering the highly repetitive structure of the NRPS modules, such systems are poorly assembled with only short reads. We also observed that MegMviri4 and VGEX01 share extremely similar CDPS systems (figure 7). Overall, according to blastp, the CDPS, NRPS and RiPP systems were most like those found in the two existing ‘*Ca*. Megaira’ genomes, MegCarteria and MegNEIS296 (Supplementary table S10).

**Figure 7.**
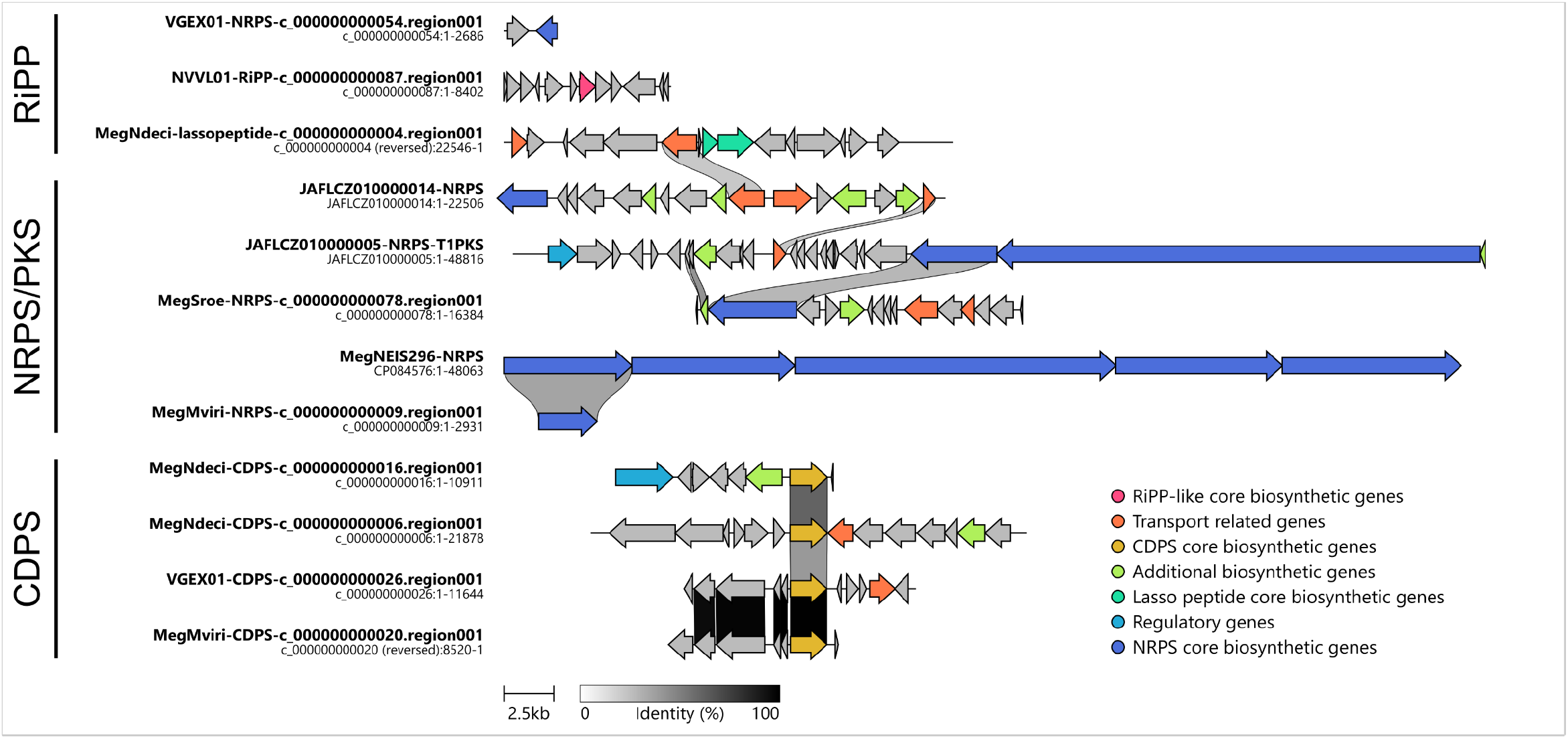
*Clinker similarity diagram of RiPP, NRPS and CDPS gene regions found across* Megaira *by antiSMASH. Similarities between genes are indicated with grey shaded links between genes, and colours represent the types of genes present as found by antiSMASH. Rows are ordered by best overall similarity according to clinker defaults. A fully interactive clinker diagram with more details on each gene function can be found in supplementary data:* https://doi.org/10.6084/m9.figshare.20424894.v1.

A mostly complete flagellar apparatus was also identified in JAFLDA01 (Supplementary table S5 and Supplementary figure 3). Partial flagella pathways are also annotated in the genomes NVVL01 and RXKF01 (Supplementary table S5). Aside these, Megaira strains all carry Sec and Tat systems for translocation of proteins to the periplasmic space, alongside one or more Type IV secretion systems (Supplementary S4).

We examined the Megaira genomes for ORFs with three classes of motif associated with protein-protein interaction considered important in symbiont-host interactions: ankyrin repeat domains, Tetratricopeptide repeats, and Leucine Rich Repeats. These genes sets were not generally common across ‘*Ca*. Megaira’ (Table 2). However, the MegNDeciBESST genome was notably enriched, including 15 ORFs carrying ankyrin repeats, 20 with predicted tetratricopeptide repeat motifs and four with leucine rich repeat genes. Two other strains NVVL01 and JAFLCZ01 have modestly increased complement of ORFs in this class (Table 2).

**Table 2.** *Number of ORFs in* Megaira *genomes containing putative protein-protein interaction domains as recognised in pfam searches*.

|  | ORF feature |  |  |
| --- | --- | --- | --- |
|  | Ankyrin domains | Tetratricopeptide repeats | Leucine Rich Repeats |
| **MegNdeciBESST** | 15 | 20 | 4 |
| **MegSroe9** | 1 | 0 | 1 |
| **MegCarteria** | 1 | 1 | 1 |
| **MegMviri4** | 1 | 3 | 1 |
| **MegNEIS296** | 1 | 4 | 1 |
| **NVVL01** | 9 | 7 | 2 |
| **JAFLDA01** | 4 | 3 | 0 |
| **JAFLCZ01** | 5 | 4 | 7 |
| **JAJTEK01** | 2 | 3 | 1 |
| **VGEX01** | 4 | 4 | 2 |

**Supplementary figure 3.**
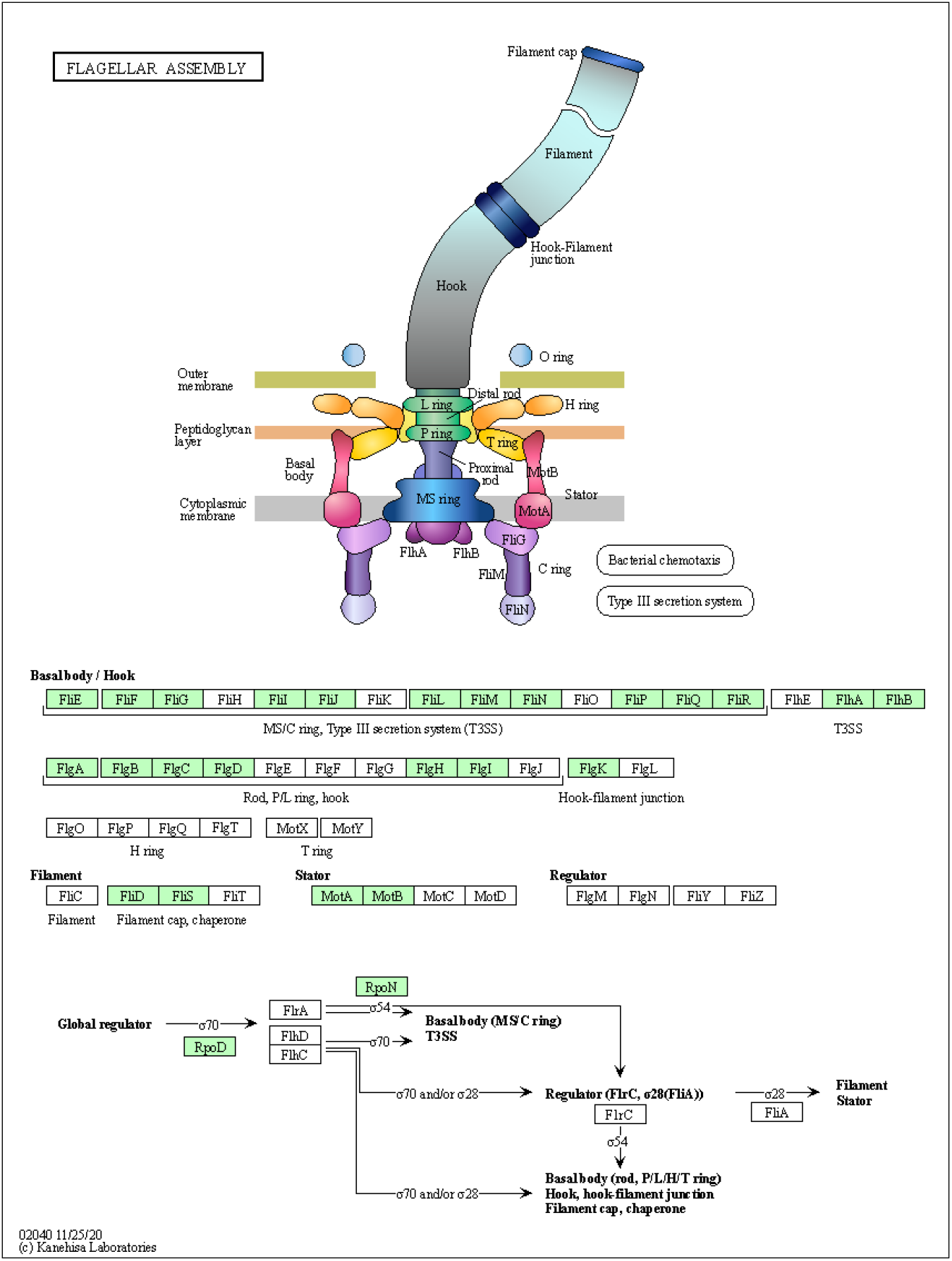
Flagella apparatus for JAFLDA01. Green highlighted boxes indicated genes found.

## Discussion

Advances in metagenomic methods and data-mining techniques are enriching our understanding of microbial symbiont diversity. The genus ‘*Ca*. Megaira’ represents a common and hyperdiverse clade of intracellular symbionts associated with microeukaryotes and algae. Using a metagenomic approach, we have assembled draft genomes for four ‘*Ca*. Megaira’ species. One of these genomes was assembled into a single scaffold using mate pair reads. In addition, we identified 14 previously existing MAGs in GenBank derived from previous environmental metagenome projects [52–63]. Of these, 5 can be considered high quality (>90% complete, <10% contamination).

Our data indicate ‘*Ca*. Megaira’ is diverse enough to be considered its own family within Rickettsiales. The available genomes for previously recognised Clades of ‘*Ca*. Megaira’ share AAI similarity below 65% as well as very low synteny between the two most complete genomes JAFLDA01, ‘*Ca*. Megaira’ Clade E and ‘*Ca*. Megaira’ Clade A from *Mesostigma viride*, (Figure 1–4, and Supplementary figure 2). In addition, NVVL01, while firmly positioned within ‘*Ca*. Megaira’, has an enormous number of unique and unclassified gene clusters that exceed all other genomes described here; this novelty indicates a potentially enormous scope for further genomic diversity within the ‘*Ca*. Megaira’ clades. Our data also indicate a new species group within the current Clade A (Figure 1 and 3). Overall, the analysis of our current and limited genomic data suggest that ‘Ca. Megaira’ lineage consist of at least 6 genus-level clades and 9 species.

Nevertheless, our understanding of ‘*Ca*. Megaira’ genomic diversity remains limited, as we are unable to consolidate the taxonomy for single genomes that fall outside the main clades or that lack 16S rRNA resulting from metagenomic assembly [64]. As such, whilst our data indicates taxonomic revision is necessary, we have chosen not to challenge current levels of taxonomic classification to avoid confusion while our knowledge of this family of bacteria is still relatively small. Instead, we encourage future studies to diversify the markers that they use for identifying ‘*Ca*. Megaira’ beyond 16S rRNA, and to obtain greater genomic information, particularly beyond clade A strains, to allow firm resolution of ‘*Ca*. Megaira’ genomic diversity to allow this revision.

All ‘*Ca*. Megaira’ genomes obtained have similar predicted metabolic potential which match the two currently available genomes for this group (Figure 5). Apart from some partial B vitamin pathways in JAFLDA01 there is little evidence of capacity for vitamin dependent nutritional symbioses in these bacteria (Figure 5 and Supplementary table S6). Most Clade A strains encode for a large number of proteins related to terpenoid and polyketide pathways (Figures 5 and 6). These are known to be associated with plant-mycorrhizal and sponge-alphaproteobacteria defensive symbioses [65]. Terpenes are also produced by algae for defence systems, and some red algae appear to be reliant on bacteria-like terpene pathways to do so [66]. Terpenoids and polyketides can also increase host tolerance to various environmental stresses including pathogenic bacteria and heavy metal pollution [65, 67]. In addition, MegNdeciBESST, which was recovered from a brown alga genome project, has a complete dTDP-L-Rhamnose biosynthesis pathway which can be associated with establishing symbiosis in plants [68, 69]. Therefore, it is possible that ‘*Ca*. Megaira’ form a type of defensive symbiosis with their hosts. However, these terpenoids could alternatively be part of establishing infection in the host algae, rather than a defensive symbiosis because bacteria use them to produce components of their cell walls [70].

The presence of systems predicted to synthesize secondary metabolites (NRPS, CDPS and RiPPs, including a lasso peptide) provide additional evidence that ‘*Ca*. Megaira’ could be involved in protective symbiosis, or a toxin-antitoxin system which can be associated with reproductive manipulation [71]. These peptide groups cover a wide variety of bacterial secondary metabolites, many of which are associated with antimicrobial, antifungal, antiviral, and antibiotic properties [72, 73]; lasso peptides additionally show very high level of tolerance to environmental extremes of temperature and pH [74]. Alternatively, the products of these systems could be actively harmful to the host as some of these molecules, like the RiPP nostocyclamide, have been shown to have anti-algal properties [75]. It is currently unknown if these systems are functional or how the products might affect their hosts. However, they do seem to be common in ‘*Ca*. Megaira’ as they are present in six of the eight genomes examined here.

Some intracellular symbionts deploy an array of proteins which interact with host proteins to modify host cellular systems and establish symbiosis. The most widely recognised of these is the expansion of genes carrying ankyrin domains in *Wolbachia* [76, 77].

MegNDeciBESST is evolutionary distant from other ‘*Ca*. Megaira’ and has a clearly expanded repertoire of genes encoding ankyrin domains, tetratricopeptide domains and leucine rich repeat domains which are associated with protein-protein interaction. This distinction likely makes the molecular basis of its symbioses distinct from that of the other strains. The MegNDeciBESST genome is particularly interesting, as it indicates the expansion of potential effectors functioning through protein-protein interaction that is observed in *Wolbachia* is not unique and has independently evolved in other intracellular symbionts. This aspect of the MegNDeciBESST genome also supports the biological diversity of symbiosis that exists within the current clade ‘*Ca*. Megaira’.

We also found evidence for a complete flagellar apparatus in a clade D Megaira, JAFLDA01. Although Rickettsiaceae do not typically have flagella, microscopy results suggested the presence of a putative flagellar structure in ‘*Ca*. Megaira’ venefica [19], a member of ‘*Ca*. Megaira’ clade C. The apparatus is also present in a few related genera such as *Ca*. Trichorickettsia and *Ca*. Gigarickettsia [78]. The presence of flagellar genes in a deep member of the Rickettsiaceae [79] further suggest that a flagellar assembly apparatus might have been an ancestral feature of Rickettsiaceae which was subsequently lost from most of the lineages. We do not know if these pathways are functional, but it is notable that complete or near complete sets of these genes are found in several ‘*Ca*. Megaira’ species, while the majority of Rickettsiaceae lack them entirely.

In conclusion, ‘Ca. Megaira’ is emerging as a diverse, cosmopolitan clade of bacteria that often form symbioses with a variety of ciliate, micro- and macro-algae. It is commonly found in aquatic metagenomes [19] and is likely associated with many other microeukaryotes. We assembled 4 new draft genomes and identify 14 existing environmental MAGs. It is still unclear how these bacteria interact with their hosts, but the presence of partial terpene pathways, alongside the occurrence of various ORFs, NRPS, CDPS and RiPPs across ‘Ca. Megaira’ could point towards defensive symbioses. We do not believe that the current taxonomy of ‘Ca. Megaira’ sufficiently describes the diversity we observe here. However, further investigation is needed to fully consolidate the identity of genomes lacking 16S rRNA and increasing genome representation to avoid clades being represented by a single genome clade. Once this is complete, the diversity and biology of this hyper diverse group can be established with greater power.

## Acknowledgements

With thanks to Dr Ying Yan and Chao Li from the Ocean University of China, Qingdao for allowing us use of their *Stentor roeselii* SRA data, bioproject PRJNA507905.

